# It’s a Trap?! Escape from an ancient, ancestral sex chromosome system and implication of *Foxl2* as the putative primary sex determining gene in a lizard (Anguimorpha; Shinisauridae)

**DOI:** 10.1101/2023.07.05.547803

**Authors:** Brendan J. Pinto, Stuart V. Nielsen, Kathryn A. Sullivan, Ashmika Behere, Shannon E. Keating, Mona van Schingen-Khan, Truong Quang Nguyen, Thomas Ziegler, Jennifer Pramuk, Melissa A. Wilson, Tony Gamble

**Affiliations:** School of Life Sciences, Arizona State University, Tempe, AZ USA; Center for Evolution and Medicine, Arizona State University, Tempe, AZ USA; Department of Zoology, Milwaukee Public Museum, Milwaukee, WI USA; Department of Biological Sciences, Museum of Life Sciences, Louisiana State University-Shreveport, Shreveport, LA USA; Florida Museum of Natural History, University of Florida, Gainesville, FL USA; Department of Biological Sciences, Marquette University, Milwaukee WI USA; Federal Agency for Nature Conservation, CITES Scientific Authority, Konstantinstraße 110, 53179 Bonn, Germany; Institute of Ecology and Biological Resources, Vietnam Academy of Science and Technology, 18 Hoang Quoc Viet Road, Hanoi 10072, Vietnam; Graduate University of Science and Technology, Vietnam Academy of Science and Technology, 18 Hoang Quoc Viet, Cau Giay, Hanoi 10072, Vietnam; Cologne Zoo, Riehler Straße 173, 50735 Cologne, Germany; Department of Biology, Institute of Zoology, University of Cologne, Zülpicher Straße 47b, 50674 Cologne, Germany; Center for Mechanisms of Evolution, Biodesign Institute, Tempe, AZ USA; Bell Museum of Natural History, University of Minnesota, St Paul, MN USA

## Abstract

Although sex determination is ubiquitous in vertebrates, mechanisms of sex determination vary from environmentally-to genetically-influenced. In vertebrates, genetic sex determination is typically accomplished with sex chromosomes. Groups like mammals maintain conserved sex chromosome systems, while sex chromosomes in most vertebrate clades aren’t conserved across similar evolutionary timescales. One group inferred to have an evolutionarily stable mode of sex determination is Anguimorpha, a clade of charismatic taxa including: monitor lizards, Gila monsters, and crocodile lizards. The common ancestor of extant anguimorphs possessed a ZW system that has been retained across the clade. However, the sex chromosome system in the endangered, monotypic family of crocodile lizards (Shinisauridae) has remained elusive. Here, we analyze genomic data to demonstrate that *Shinisaurus* has replaced the ancestral anguimorph ZW system on LG7 chromosome with a novel ZW system on LG3. The linkage group LG3 corresponds to chromosome 9 in chicken, and this is the first documented use of this syntenic block as a sex chromosome in amniotes. Additionally, this ∼1Mb region harbors approximately 10 genes, including a duplication of the sex-determining transcription factor, *Foxl2*—critical for the determination and maintenance of sexual differentiation in vertebrates, and thus a putative primary sex determining gene for *Shinisaurus*.

## Introduction

The evolution of sex determination in vertebrates is impressive in its ability to combine a highly conserved developmental network that can be initiated by quite distinct molecular mechanisms in different species (Bachtrog et al. 2014; Graves, 2008). In vertebrates, sex is commonly determined via either environmental and/or genetic cues at critical points in development. In vertebrate groups that use genetic mechanisms, the most common mechanism is sex chromosomes; either a male or female heterogametic system where the male or female inherits the sex-limited (Y or W) chromosome, respectively (Bachtrog et al. 2014; Gamble et al. 2015). Sex chromosomes have been traditionally identified by comparing male and female karyotypes under the light microscope. The presence of morphological differences between the X and Y (or Z and W) chromosomes (i.e. heteromorphic sex chromosomes) identify a species’ sex chromosome system (Stevens, 1905; Bull, 1983). However, many species possess sex chromosomes that cannot be identified via light microscopy because the X and Y (or Z and W) are not morphologically distinguishable from each other (i.e. homomorphic sex chromosomes). Other methods must be employed, such as advanced cytogenetic techniques or high-throughput DNA sequencing technologies, to identify sex chromosome systems in these taxa (Gamble and Zarkower, 2014; Gamble et al. 2017; Pinto et al. 2022).

Squamate reptiles (lizards and snakes) demonstrate high variability in modes of sex determination: where some clades have conserved, often heteromorphic, sex chromosomes, while others display extraordinary lability in their modes of sex determination and a high incidence of homomorphic sex chromosomes (Gamble et al. 2015, Kratochvíl et al. 2021; Augstenová et al. 2021a). One hypothesis of sex chromosome evolution is that ancient, degenerated sex chromosome systems may act as an “evolutionary trap”, where the existence of highly differentiated (i.e. heteromorphic) sex chromosomes preclude transitions to other sex-determining systems (Bull 1983; Bull and Charnov, 1985; Pokorná and Kratochvíl, 2009). The stability of old sex chromosome systems in mammals, birds, caenophidian snakes, and others, provides anecdotal support for this hypothesis (Bull and Charnov, 1985; Pokorná and Kratochvíl, 2009; Gamble et al. 2015). As more and more sex chromosome transitions are identified, it remains unclear whether all ancient sex chromosome systems are destined to become traps, but examples of taxa transitioning away from ancient, degenerated sex chromosome systems are rare in amniotes (Acosta et al. 2019; Nielsen et al. 2019; Rovatsos et al. 2019a). Previous phylogenetic studies have supported the trap hypothesis in squamates (Pokorná and Kratochvíl, 2009; Gamble et al. 2015), but also suffered from incomplete taxonomic sampling, which might have biased the conclusions. In other words, testing this hypothesis is contingent upon having sufficient data necessary to identify transitions away from an ancient sex chromosome system, which typically requires (1) a reference genome to coordinate linkage groups (which are rare in squamates; Pinto et al. 2023), (2) genome-scale data from both sexes (e.g. Vicoso et al. 2013; Gamble et al. 2015; Pinto et al. 2022), and (3) a robust phylogenetic hypothesis to establish relationships within the focal taxa (Nielsen et al. 2019). Thus, the burden of proof is higher for identifying escapees from these ancient sex chromosome systems, which may be responsible for the dearth of examples and the previous lack of conclusive examinations of the evolutionary trap hypothesis. Future identification of additional escapees will permit more conclusive analyses of whether or not ancient sex chromosome systems truly act as evolutionary traps across a broader phylogenetic scale.

The sex chromosomes of the infraorder Anguimorpha (lizards including monitor lizards, Gila monsters, alligator lizards, and their allies) have long been a topic of interest, likely resulting from the paucity of genetic and cytogenetic data for this group. In recent years, advanced cytogenetic techniques (FISH) have facilitated karyotypic analysis and identification of ZW sex chromosomes in the Gila monster (*Heloderma suspectum*; Pokorná et al. 2014) and Komodo dragon (*Varanus komodoensis*; Pokorná et al. 2016) leading to expanded interest in studying chromosome evolution in this enigmatic group. More recently, RNAseq and qPCR analysis, in conjunction with draft genomes of these same two anguimorph species (Gila monster; Webster et al. 2023, and Komodo dragon; Lind et al. 2019), have provided some additional insights into this system (Rovatsos et al. 2019b). Namely, the homology of the heteromorphic ZW systems in the anguimorph genera *Abronia*, *Heloderma*, and *Varanus* (Rovatsos et al. 2019b; Webster et al. 2023). The presence of a ZW sex chromosome on the same linkage group—syntenic with chromosome 28 in the chicken genome—in these three genera, spanning the phylogenetic breadth of extant Anguimorpha, is strong evidence that this is the ancestral sex chromosome system in the clade. Ancient sex chromosome systems, like those ancestral to anguimorphs (115–180 million years old), fit the criteria that should render them as an evolutionary trap (Pokorná and Kratochvíl, 2009; Rovatsos et al. 2019b). However, the sex chromosomes of many anguimorph taxa remain unknown, including the monotypic family Shinisauridae, which is nested within the anguimorph phylogeny (Figure 1).

**Figure 1:**
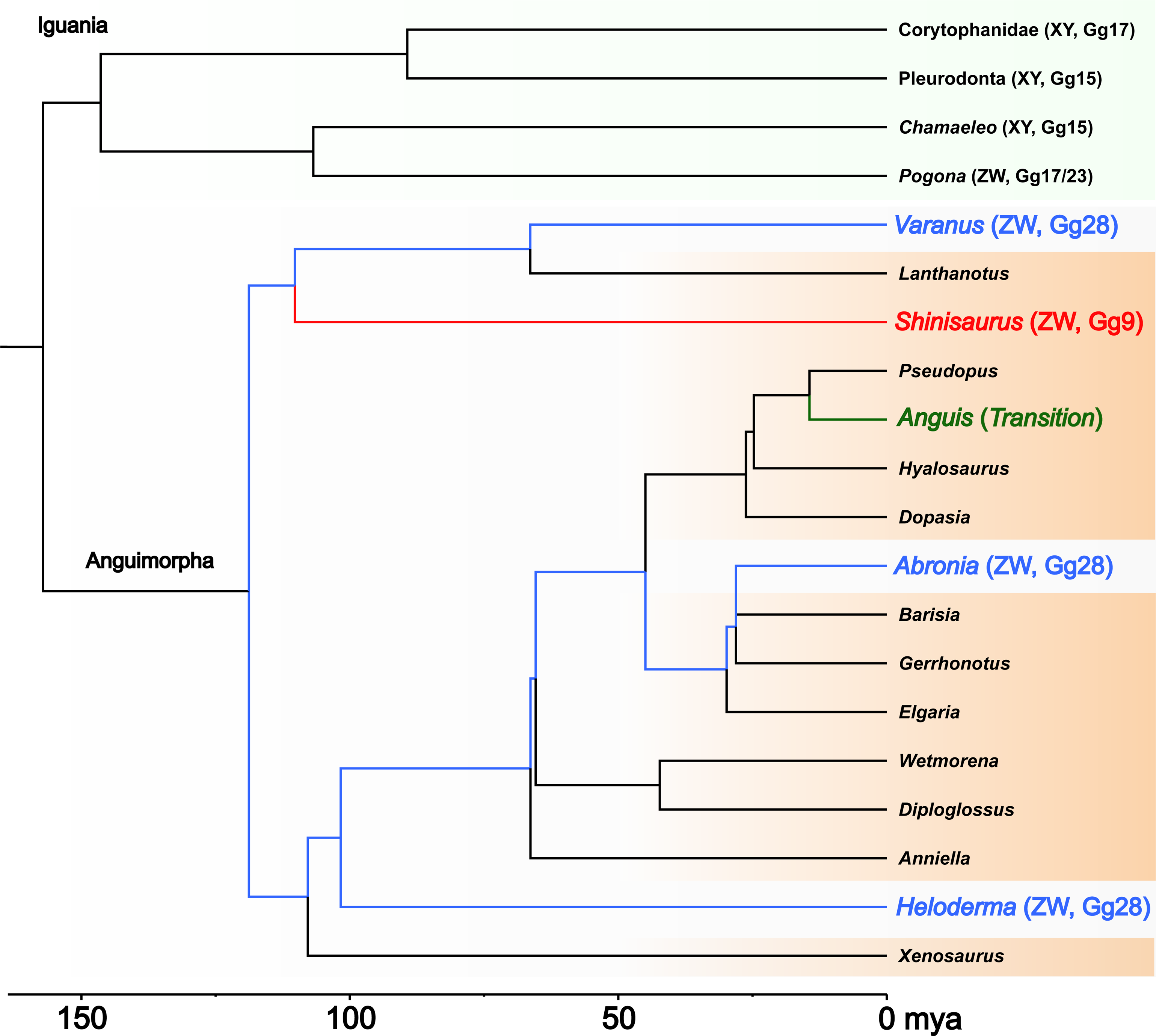
Summary of current anguimorph sex chromosome knowledge summarized from Rovatsos et al. (2019b) indicated by blue and green tips/branches); information identified in this study indicated by red tips/branches and what remains unknown across the phylogeny indicated by black tips/branches. Phylogeny from TimeTree using a representative species from each clade (Kumar et al. 2017) and visualized using Figtree [v1.4.4] (http://tree.bio.ed.ac.uk/software/figtree/). Of note, “pleurodonts” represents “non-corytophanid pleurodonts” and “Gg” stands for chicken (*Gallus gallus*) linkage group.

The crocodile lizard (*Shinisaurus crocodilurus*) is the sole living member of the family Shinisauridae and native to small disjunct regions of southeastern China and northern Vietnam (Le and Ziegler, 2003; Huang et al. 2008; Nguyen et al. 2014). It is one of the rarest lizard species in the world and is listed as Endangered in the IUCN Red List (Nguyen et al. 2014). Due to anthropogenic pressures, populations have experienced severe declines in recent years with less than 1000 individuals in the wild in China and less than 100 adults in Vietnam (Huang et al. 2008; van Schingen et al. 2014a). They are semi-aquatic habitat specialists and depend upon clean streams in broadleaf evergreen forest (Ning et al. 2006; van Schingen et al. 2016a) and their restricted ecological niche is predicted to all but disappear due to climate change by the end of this century (Li et al. 2013; van Schingen et al. 2014a; see also van Schingen-Khan et al. 2022). Habitat destruction threatens remaining populations, as well as overcollection for food and the international pet trade (Huang et al. 2008; van Schingen et al. 2014b; van Schingen et al. 2016a). Although still recognized as a single species, there exist multiple conservation units, with *S. crocodilurus vietnamensis* from Vietnam and the nominal subspecies from China consisting of several distinct lineages (van Schingen et al. 2016b; Ngo et al. 2020; Nguyen et al. 2022). Crocodile lizards do not have a clear sexual dimorphism. While morphological traits, such as coloration or body morphometry, may provide some indication of the sex, it remains difficult for most people to identify the sex of individuals (van Schingen et al. 2016b). Relevant to the present study, examination of male and female *S. crocodilurus* karyotypes have revealed no heteromorphic sex chromosomes (Zhang et al. 1996; Augstenová et al. 2021b). To identify sex chromosomes in this species, we analyzed whole-genome re-sequencing data for approximately 50 sexed, individual crocodile lizards (Xie et al. 2022) using whole-genome re-sequencing to show that the sex determining system in *S. crocodilurus* is a novel ZW system that has eluded previous analyses, at least in part, due to the small size (<1Mb) of its sex determining region (SDR).

## Methods

### WGS analysis

We downloaded low-coverage whole genome Illumina resequencing (WGS) reads from NCBI SRA for multiple male and female individuals (see *Data Availability* for accessions). We constructed a Snakemake [v6.10.0] (Mölder et al. 2021) workflow in an isolated conda environment [v4.11.0] (https://docs.anaconda.com/) containing relevant packages: BBmap [v38.93] (Bushnell, 2014), FastQC [v0.11.9] (Andrews, 2010), Freebayes [v1.3.5] (Garrison and Marth, 2012), GFF utilities [v0.10.1] (Pertea and Pertea, 2020), Minimap2 [v2.22] (Li, 2018), Mosdepth [v0.3.2] (Pedersen and Quinlan, 2018), MultiQC [v1.11] (Ewels et al. 2016), Parallel [v20211022] (Tange, 2018), pixy [v1.2.5.beta1] (Korunes and Samuk, 2021), RTGTools [v3.12.1] (Cleary et al. 2015), Sambamba [v0.8.1] (Tarasov et al. 2015), Samtools [v1.12] (Li et al. 2009), seqkit [v0.11.0] (Shen et al. 2016), STACKS [v2.6.0] (Catchen et al. 2013), and Trim Galore! [v0.6.7] (https://doi.org/10.5281/zenodo.5127899). To process the raw sequencing data, we trimmed adapters and low-quality regions using Trim Galore!, then removed PCR duplicates using BBmap. Quality assessment using FastQC and MultiQC was conducted at each step, and we subsequently removed samples with fewer than 5 million PE reads after filtering PCR duplicates. The final WGS dataset possessed 50 sexed samples (27 male and 23 female individuals) sourced from China and Vietnam. We proceeded to map reads for each individual to the female reference genome (Xie et al. 2022) with minimap2 and calculated read depth and read mapping statistics using mosdepth and samtools, respectively. Then, we generated an all-sites VCF file with freebayes-parallel. Lastly, we calculated Weir and Cockerham (1984) F_ST_ between males and females and nucleotide diversity statistics using pixy at 500kb resolution and, for LG3 only, also at 100kb resolution.

### *Validation of the putative ZW system in* Shinisaurus crocodilurus

Male vs. female F_ST_ values are agnostic to which sex is heterogametic (i.e. XY vs. ZW). Therefore, we generated a dataset of ‘*in silico* poolseq’ reads by subsampling each WGS sample to 10 million paired reads (20 million total reads per sample) using seqkit and combined into male and female pools. We analyzed the pools using Pooled Sequencing Analyses for Sex Signal [PSASS; v3.1.0] (https://doi.org/10.5281/zenodo.3702337). We then generated PCR primers targeting the annotated version of *Foxl2*’s second exon [FOXL2-ex2-F2 5’ – CAGAGCTCGTCCCATTCACTT – 3’ and FOXL2-ex2-R2 5’ – GAGAGATGTACCACCGGGAG – 3’] and sequenced the resultant amplicon using Sanger sequencing (Psomagen). Individuals used in Sanger sequencing are detailed in Supplemental Table 1.

### Genome Annotation

We used previously lifted over annotations (Pinto et al. 2023; https://doi.org/10.6084/m9.figshare.20201099.v1) via Liftoff [v1.6.3] (Shumate and Salzberg, 2021) from the draft genome of a male *S. crocodilurus* (Gao et al. 2017) to the new, unannotated female reference genome (Xie et al. 2022; GCA_021292165.1). We pulled coding transcripts from the genome using GFF Utilities. We used the 10 genes within the putative ∼1Mb SDR on LG3 to perform a high-stringency tBLASTx query (Altschul et al. 1990) to the chicken genome on Ensembl (Howe et al. 2020) with a word size of 3, maximum of 10 hits, e-value cutoff of 1e^−50^, using BLOSUM62 scoring matrix. These queries received hits on 7 of the 10 total genes (Table 1).

**Table 1:**
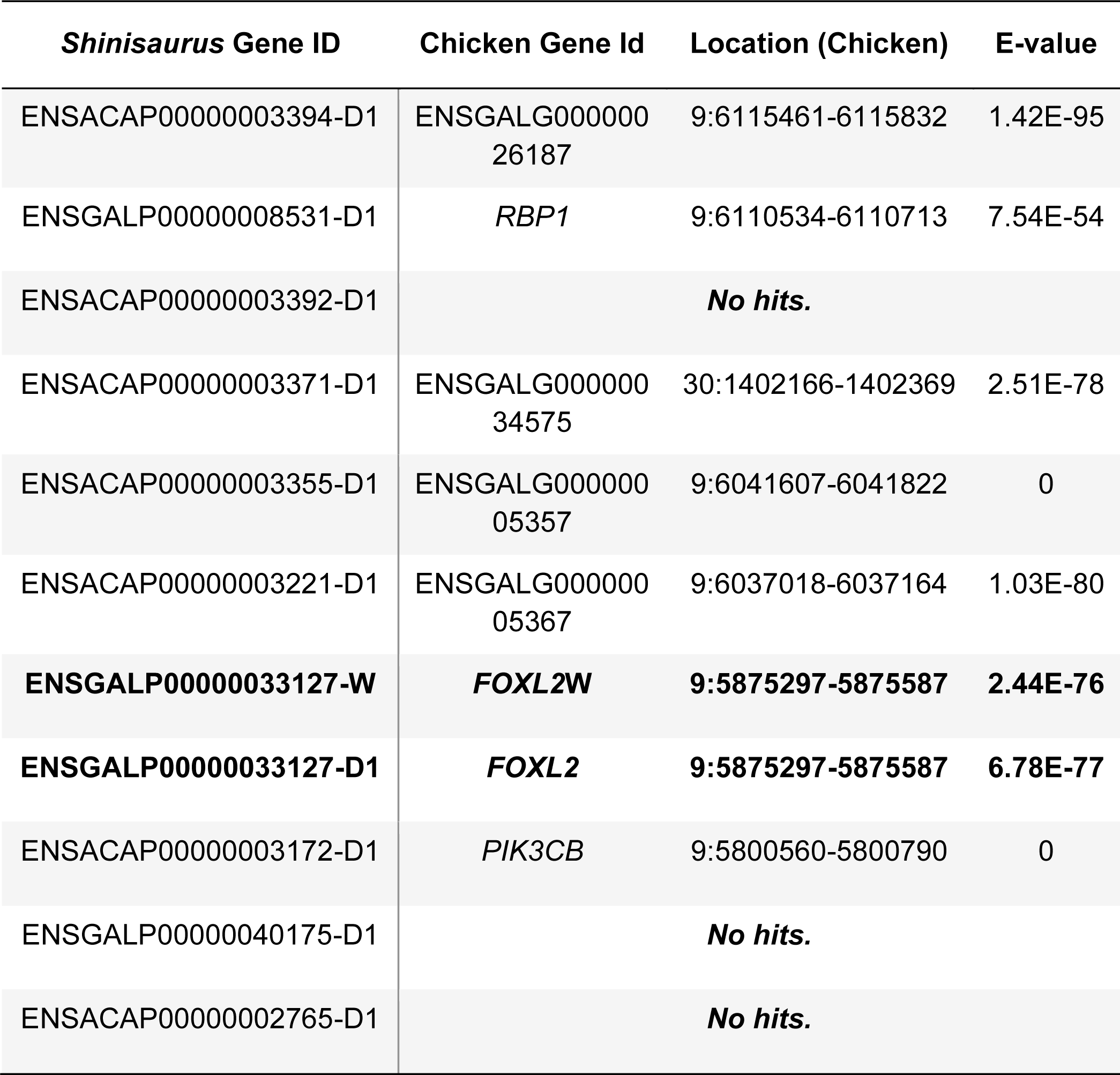
Top tBLASTx hits in chicken for the CDS of each gene present in the *Shinisaurus* 900kb-SDR. The duplicated *Foxl2* copy is dubbed ENSGALP00000033127-W.

| *Shinisaurus* Gene ID | Chicken Gene Id | Location (Chicken) | E-value |
| --- | --- | --- | --- |
| ENSACAP00000003394-D1 | ENSGALG00000026187 | 9:6115461-6115832 | 1.42E-95 |
| ENSGALP00000008531-D1 | *RBP1* | 9:6110534-6110713 | 7.54E-54 |
| ENSACAP00000003392-D1 | **No hits.** |  |  |
| ENSACAP00000003371-D1 | ENSGALG00000034575 | 30:1402166-1402369 | 2.51E-78 |
| ENSACAP00000003355-D1 | ENSGALG00000005357 | 9:6041607-6041822 | 0 |
| ENSACAP00000003221-D1 | ENSGALG00000005367 | 9:6037018-6037164 | 1.03E-80 |
| **ENSGALP00000033127-W** | ***FOXL2W*** | **9:5875297-5875587** | **2.44E-76** |
| **ENSGALP00000033127-D1** | ***FOXL2*** | **9:5875297-5875587** | **6.78E-77** |
| ENSACAP00000003172-D1 | *PIK3CB* | 9:5800560-5800790 | 0 |
| ENSGALP00000040175-D1 | **No hits.** |  |  |
| ENSACAP00000002765-D1 | **No hits.** |  |  |

## Results

Across WGS experiments, read mapping efficiency ranged from 80.60% (for SRR5019740) to 99.40% (for SRR14583318). After variant calling, the WGS dataset contained 6,202,005 biallelic variants (see Data Availability section for additional VCF statistics). We identified a region of high F_ST_ between males and females on linkage group 3 (LG3; Figure 2), however, comparing M/F F_ST_ values does not necessarily diagnose which sex is heterogametic (i.e. XY vs. ZW). Therefore, we composed a dataset of ‘*in silico* poolseq’ reads to identify an excess of female-associated SNPs aligning to the previously identified region of high M/F F_ST_ (Supplemental Figure 1). Taken together, these data suggest that *S. crocodilurus* possesses a female heterogametic system (ZW) with an SDR located in a ∼900kb region on LG3.

**Figure 2:**
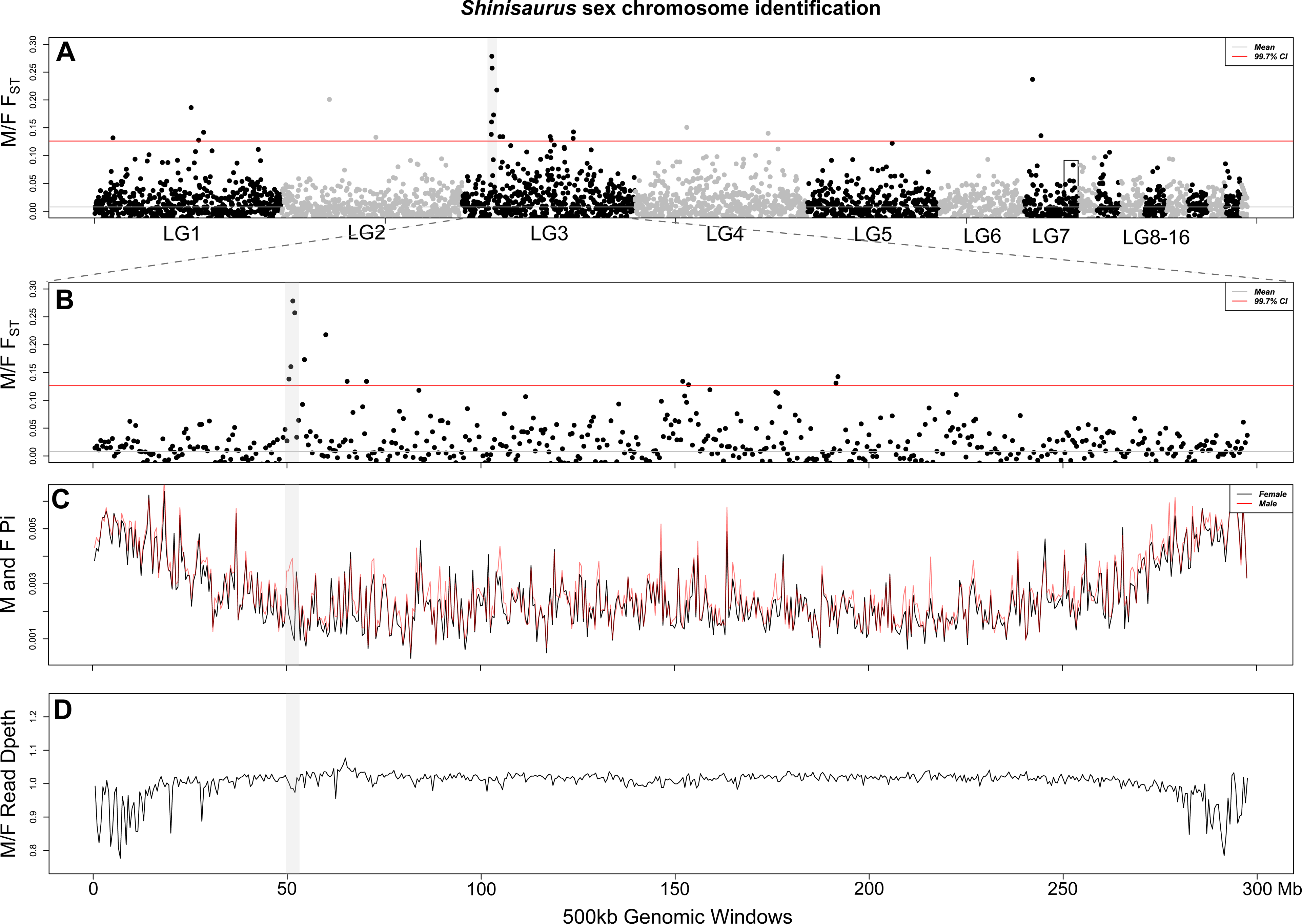
Identification of the ZW sex chromosome system in *Shinisaurus crocodilurus*. (A) Whole genome F_ST_ scan with a clear peak in a ∼1Mb region on LG3. The square block on LG7 is syntenic with the sex-determining region in *Varanus* and *Heloderma* (Webster et al. 2023). (B) Isolation and magnification of LG3 F_ST_ peak. (C) Modest increase in male, relative to female, nucleotide diversity and (D) decrease in male/female read depth in the region corresponding to the F_ST_ peak on LG3.

Upon further investigation of the SDR, we identified a total of 10 genes annotated within this region of high F_ST_ and an excess of female-specific SNPs. To better characterize these genes, we BLAST-ed each to the chicken genome. We recovered high-quality BLAST hits for seven of the 10 annotated *S. crocodilurus* SDR genes in chicken (Table 1). Six out of the seven queries hit genes located on chicken chromosome 9, while the other landed on a chicken chromosome 30 (Table 1). In our poolseq analysis, one of these genes possessed half the read depth in females relative to males (Supplemental Table 2) and, upon closer inspection, we identified a duplicated, unannotated copy of that gene Forkhead Box L2 (*Foxl2*), located approximately 70kb upstream—with 99% sequence identity, also located within the putative SDR. We included this *Foxl2* copy in a BLAST search against chicken, where it was again identified as a *Foxl2* homolog (Table 1). We also BLAST-ed *Foxl2* to the earlier male *S. crocodilurus* draft genome (Gao et al. 2017) and found only a single copy of *Foxl2* in this genome matching one copy in the updated reference genome with 100% sequence identity, consistent with both (1) the duplicated version being W-specific and (2) the female reference being chimeric for Z and W alleles (Xie et al. 2022). Lastly, we generated a gene tree using *Foxl2* copies from across reptiles to confirm its duplicated origination was within Shinisauridae (Supplemental Figure 2). Thus, in the chimeric female reference genome, this putative W-linked *Foxl2* copy was located approximately 70kb upstream of the annotated Z-linked copy of *Foxl2* on the other side of an assembly gap.

The WGS data used in the *in silico* PoolSeq analysis were restricted to only individuals from Chinese populations to reduce the influence of population-specific demographic processes (Xie et al. 2022). To include the less-numerous Vietnam samples, we generated PCR primers for a segment of *Foxl2*’s second exon and Sanger sequenced multiple females (Vietnam) and males (China and Vietnam) (Supplemental Figure 3). We identified one SNP in the female Vietnamese samples in this region and tested its association with sex using Fisher’s exact test (p-value = 0.0003***). Thus, the ZW SDR containing *Foxl2* appears to be conserved between populations of *S. crocodilurus* from both China and Vietnam.

## Discussion

### Escaping the “Evolutionary Trap”

An open question within sex chromosome evolution is whether ancient, degenerated sex chromosomes act as evolutionary traps (Pokorná and Kratochvíl, 2009; Nielsen et al. 2019; Pinto et al. 2023). The most recent common ancestor of extant anguimorphs is thought to have possessed a ZW system on the linkage group syntenic with chicken chromosome 28, which is located on the distal region of LG7 in in the *S. crocodilurus* reference genome (Rovatsos et al. 2019b; Webster et al. 2023). The sex determining region (SDR) in *S. crocodilurus* is located on LG3, a region syntenic with chicken chromosome 9. Of note, however, at present it is difficult to assess the precise genomic coordinates and gene content of the SDR due to the chimeric nature of the reference genome assembly. To our knowledge, this is the first demonstration in a tetrapod of the syntenic region of chicken chromosome 9 being recruited in a sex determining role (Kratochvíl et al. 2021), lending further support to the idea that all chromosomes will likely be recruited into a sex determining role given thorough enough phylogenetic sampling (Graves and Peichel, 2010; Hodgkin, 2002; O’Meally et al. 2012; Pinto et al. 2022).

It is clear from these genomic data that *S. crocodilurus* possesses a distinct sex chromosome system from all other known anguimorphs. Unlike the case of Corytophanidae and other pleurodonts, where phylogenetic relationships among taxa were inconclusive (Nielsen et al. 2019), the relationship of *S. crocodilurus* to all other anguimorphs is far less divisive. Indeed, *S. crocodilurus* is well-supported as nested within Anguimorpha—either sister to Varanidae as a member of the “Paleoanguimorpha” (Burbrink et al. 2020) or as sister to a clade containing Varanidae and Lanthanotidae (Singhal et al. 2021), depending on taxonomic sampling. Thus, assuming the hypothesis that an ancient origin of the ZW sex chromosome system possessed by extant *Varanus*, *Heloderma*, and *Abronia* is correct, then *S. crocodilurus* has successfully escaped the evolutionary trap of their ancestral, degenerated sex chromosome system—a system nearly as ancient as those systems found in both mammals and birds (Rovatsos et al. 2019b; Webster et al. 2023). It is worth noting that there remains another putative escape from the ancestral anguimorph sex chromosome system in *Anguis* that has yet to be explored further (Rovatsos et al. 2019b) and more recent phylogenetic work has implicated that Corytophanidae is likely nested somewhere within other pleurodonts, rather than being sister to all other species (Burbrink et al. 2020; Singhal et al. 2021). This suggests that there are a minimum of two evolutionary escapes within Toxicofera (snakes, iguanians, and anguimorphs)—and perhaps even two within the infraorder Anguimorpha alone.

### Primary Sex Determination in Shinisauridae

In many vertebrate groups where the primary sex determiner (PSD) is known, a relatively short list of commonly-recruited PSDs have been identified (i.e. the ‘usual suspects’; Adolfi et al. 2021; Dor et al. 2019; Herpin and Schartl, 2015). Indeed, the same genes, or their paralogs, have been independently co-opted to function as the PSD in many taxa, examples including *Sox3* in placental mammals and some medaka (members of the *Oryzias celebensis* and *O. javanicus* groups); *Amh* in tilapia, northern pike, and potentially other anguimorphs (Li et al. 2015; Myosho et al. 2015; Pan et al. 2019; Rovatsos et al. 2019b; Webster et al. 2023; and see Pan et al. 2021 for recent review); and *Dmrt1* in birds, a frog (*Xenopus laevis*), tongue sole, and other medaka fish (members of the *Oryzias latipes* group) (Chen et al. 2014; Ioannidis et al. 2021; Matsuda et al. 2002; Nanda et al. 2002; Smith et al. 2009). This is the first time Forkhead Box L2 (*Foxl2*) has been implicated as a PSD in a vertebrate, although it has been predicted to be one (e.g. Ma et al. 2022).

The transcription factor, *Foxl2*, is a direct transcriptional activator of aromatase, involved in development of the ovaries and its loss in mice during embryogenesis leads to abnormal ovarian development and infertility (Fleming et al. 2010; Pannetier et al. 2006; Schmidt et al. 2004; Uda et al. 2004). After primary sex determination and sexual development have concluded, *Dmrt1* and *Foxl2* antagonize each other transcriptionally in gonadal tissue, where sustained *Dmrt1* and *Foxl2* expression is required for adult maintenance of testis and ovary tissue, respectively (Garcia-Ortiz et al. 2009; Matson et al. 2011; Uhlenhaut et al. 2009). Indeed, *Foxl2* also behaves in a dose-dependent manner in some turtle species where its overexpression at the embryonic stage can induce male-to-female sex reversal in ZZ soft-shelled turtles (*Pelodiscus sinensis*) and female differentiation in male-temperature-incubated red-eared sliders (*Trachemys scripta*) (Jin et al. 2022; Ma et al. 2022). Importantly, *Dmrt1* has been recruited to act as a primary sex determining gene in multiple taxa (Matson and Zarkower, 2012), while *Foxl2* has remained mysteriously absent from this list—with the singular putative exception being recently described in some species of bivalve mollusks (Han et al. 2022). Thus, the identification of both *Foxl2* and a duplicated *Foxl2* copy in the W-limited region of the *Shinisaurus* genome supports the expanded list of the “usual suspects” that might act as the PSD in vertebrates.

Pragmatically, the identification of a novel ZW system in *S. crocodilurus* may present an important juncture in the conservation efforts of this endangered lizard species, that are urgently needed (Nguyen et al. 2014). Body morphometrics in mature specimens may provide an indication of the sex, i.e. males tend to have a relatively larger head, relative to abdomen length than females (van Schingen et al. 2016b). However, definitive sexually dimorphic characters are lacking in the species, especially in hatchlings, juveniles, and subadults. Therefore, a molecular genetic sex test could assist in well-managed captive breeding efforts in this species (Ziegler et al. 2019). This is vital as it’s estimated only ∼1,000 individuals remain in the wild populations in China and Vietnam during the last census (Huang et al. 2008; van Schingen et al. 2016a), while loss of remaining habitats and poaching are considered ongoing. This information may play a vital role in conservation efforts of this species and should be incorporated into ongoing captive breeding work (Ziegler et al. 2019).

In conclusion, using a combination of sequencing and validation techniques we identified the elusive ZW system in the endangered crocodile lizard, *Shinisaurus crocodilurus*. This ZW system is located on LG3 and, although interpretation inherits strong reference bias (a chimeric ZW reference genome), the SDR appears to be <1Mb in size and contains approximately 10 genes. One of these genes, *Foxl2*, possesses a duplicated copy and is important in ovarian development and fertility in vertebrates. Because of its sequence conservation (either strictly age-related or via gene conversion) and possibly its proximity to the original Z copy of *Foxl2*, we hypothesize that if *Foxl2* is the PSD in this system, it may be a gene dosage-dependent mechanism, where ZW females possess three copies of *Foxl2* instead of the two copies of ZZ males. This specific hypothesis assumes that the Z copy of *Foxl2* is retained in the pseudoautosomal region of the W chromosome, however, phased Z and W sequences are needed to provide additional support to this model. The hypothetical mechanism would essentially be the inverse of the dose-dependent *Dmrt1* sex determination in birds, where a lack of *Dmrt1* on the W decreases *DMRT1* expression in females, allowing for *Foxl2* to proceed with ovarian development (Ioannidis et al. 2021; Smith et al. 2009). Here, extra gene copies of *Foxl2* increase *FOXL2* expression to downregulate *Dmrt1* expression and initiate ovarian development in the developing gonad. Thus, we provide a putative sex determining gene for the crocodile lizard (*Shinisaurus crocodilurus*) and speculate as to its potential mechanism of action in this system.

## Data Availability

The WGS data used in this study is available on NCBI, SRA accessions for WGS data are: SRR14583317, SRR14583321, SRR14583324-26, SRR14583330, SRR14583333, SRR14583340-49, SRR14583351, SRR14583353-54, SRR14583356, SRR14583360-66, SRR5019733-45, SRR14583318-20, SRR14583322-23, SRR14583331, SRR14583334-39, SRR14583346, SRR14583350, SRR14583352, SRR14583355, SRR14583357-59. Sequence data generated in this study are available on SRA under BioProject PRJNA975696, detailed in Supplemental Table 1, and code, including and VCF statistics and gene alignments, are available on GitHub: https://github.com/DrPintoThe2nd/Shinisaurus_ZW.

## Acknowledgements

The authors would like to acknowledge Research Computing at Arizona State University for providing high-performance computing and storage resources that have contributed to the research results reported within this paper (http://www.researchcomputing.asu.edu). We thank Anna Rauhaus (Cologne Zoo) for her help with the application and preparation of tissue sending and the Woodland Park Zoo for their respective assistance. Many thanks CITES Management Authority of Vietnam for issuing permits (CITES permits No. 13VN1246N/CT-KL and 16VN0920N/CT-KL).This work was funded by the Morris Animal Foundation (Study grant D19ZO-021) for their generous funding of this project (T.G.) and also supported by the National Institute of General Medical Sciences (NIGMS) of the National Institutes of Health grant R35GM124827 (M.A.W.).

