## Supplemental for "It’s a Trap?! Escape from an ancient, ancestral sex chromosome system and implication of *Foxl2* as the putative primary sex determining gene in a lizard (Anguimorpha; Shinisauridae)"

**Supplemental Figure 1:** *In silico* PoolSeq analysis using PSASS to identify sex-associated SNPs. Locations of increased female-specific SNPs coincide with high $F_{ST}$ peak in WGS M/F $F_{ST}$ analysis in Figure 2 of the main text (Supplemental Figure 1).

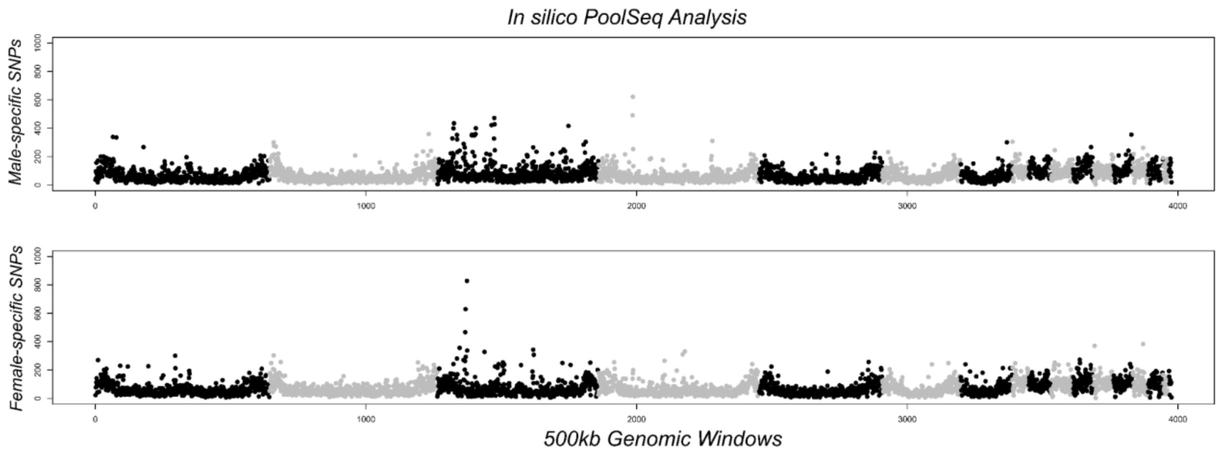

**Supplemental Figure 2:** *Foxl2* gene tree for representative sauropsid taxa demonstrating the two gametologous copies (Z and W copies) of *Foxl2* present in *Shinisaurus*. Gene tree generated using IQ-Tree [v2.2.2.3] (Nguyen et al. 2014) with 1000 UF-boot replicates and visualized using FigTree [v1.4.4] (http://tree.bio.ed.ac.uk/software/figtree/).

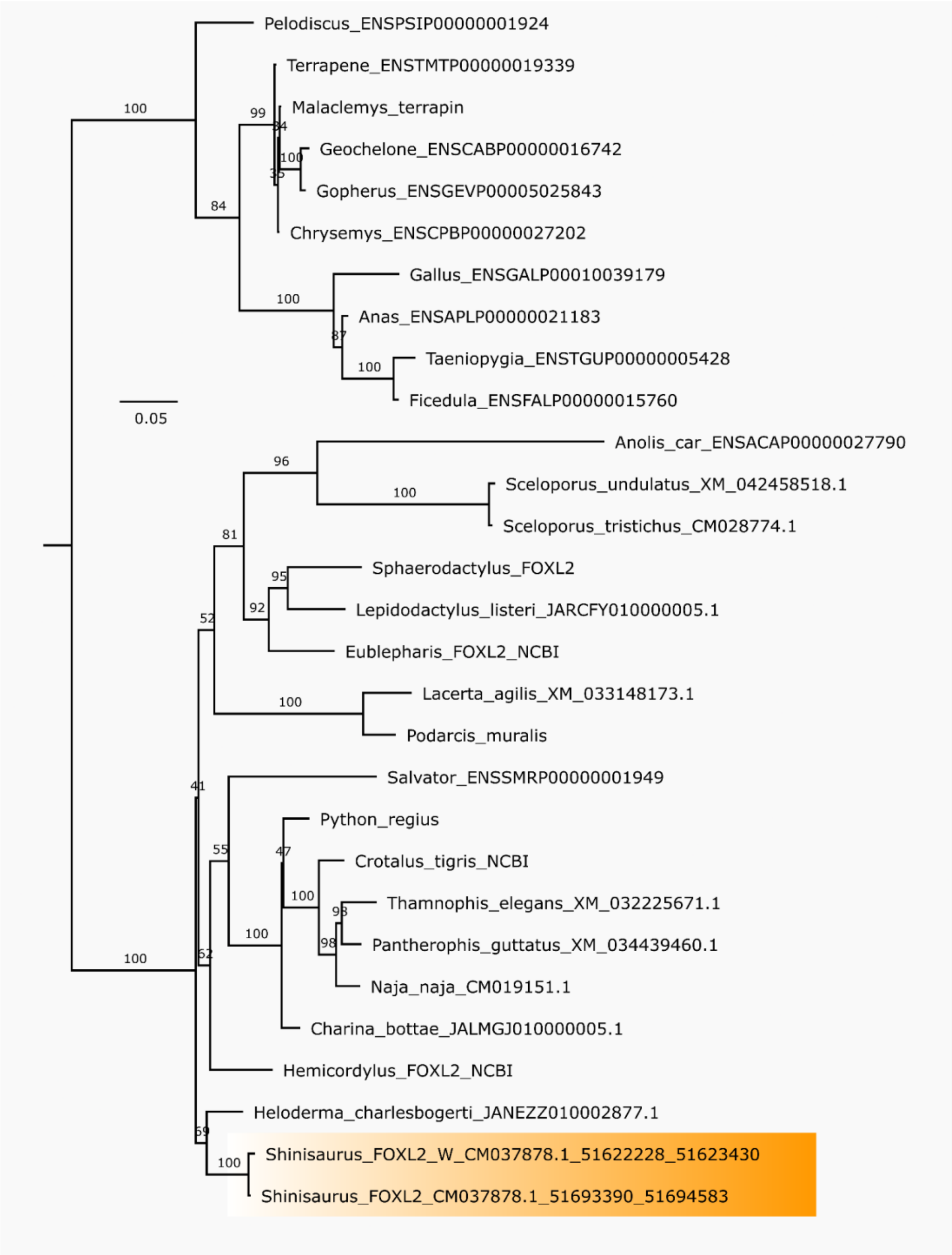

**Supplemental Figure 3:** Sanger sequencing data for *Foxl2*'s 2nd exon in multiple males and females. Here, the Z-linked copy possesses a Proline residue at the 12th AA position in the alignment, while the W-linked copy possesses an Leucine at this position. Due to sequence conservation between the two copies in females, both homozygous copies are amplified and appear as a heterozygous site in the alignment. Heterozygosity at this site is significantly associated with sex using Fisher's exact test (p-value = 0.0047***).

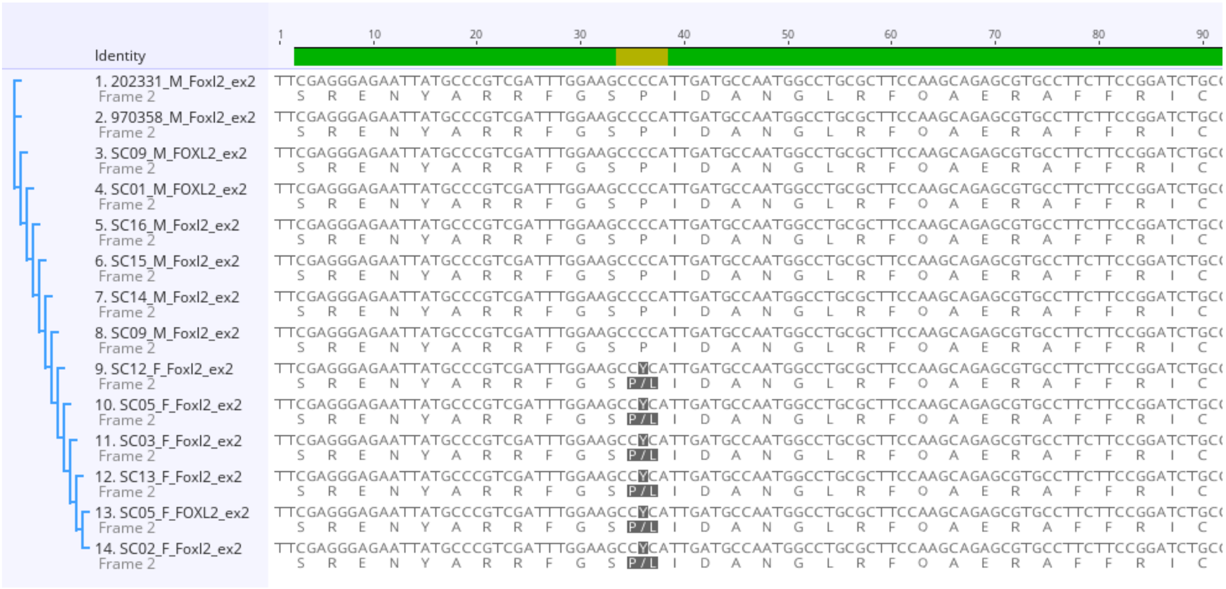

| **Supplemental Table 1:** Metadata for individuals Sanger sequenced in this study (WPZ == Woodland Park Zoo and MLSB == Me Linh Station for Biodiversity). We also generated a modest RADseq and HiC dataset that was omitted from the final manuscript but is detailed here for additional documentation that the data is publicly available for reuse. | | | | |
|---|---|---|---|---|
| Individual | Type | SRA | Sex | Sample Source |
| Sc1 | Sanger/RADseq | SRR24714874 | M | Vietnam (MLSB) |
| Sc2 | Sanger | N/A. | F | Vietnam (MLSB) |
| Sc3 | Sanger/RADseq | SRR24714873 | F | Vietnam (MLSB) |
| Sc5 | Sanger/RADseq | SRR24714872 | F | Vietnam (MLSB) |
| Sc9 | Sanger | N/A. | M | Vietnam (MLSB) |
| Sc10 | RADseq | SRR24714877 | M | Vietnam (MLSB) |
| Sc12 | Sanger/RADseq | SRR24714876 | F | Vietnam (MLSB) |
| Sc13 | Sanger | N/A. | F | Vietnam (MLSB) |
| Sc14 | Sanger | N/A. | M | Vietnam (MLSB) |
| Sc15 | Sanger | N/A. | M | Vietnam (MLSB) |
| Sc16 | Sanger/RADseq | SRR24714875 | M | Vietnam (MLSB) |
| 200322 | RADseq | SRR24714882 | F | China (WPZ) |
| 202331 | Sanger/RADseq | SRR24714881 | M | China (WPZ) |
| 205243 | RADseq | SRR24714879 | F | China (WPZ) |
| 205771 | RADseq | SRR24714878 | F | China (WPZ) |
| 205771 | HiC | SRR24714880 | F | China (WPZ) |
| 970358 | Sanger | N/A. | M | China (WPZ) |

| **Supplemental Table 2:** PSASS depth output for the annotated copy of *Foxl2*. F_depth_corr and M_depth_corr stand for corrected PoolSeq read depth for females and males, respectively. | | | | | |
|---|---|---|---|---|---|
| Contig | Start | End | ID | F_depth_corr | M_depth_corr |
| LG_3 | 51693393 | 51707080 | ENSGALP00000033127-D1 | 5 | 10 |
